# Temporal resolution reshapes dynamics of inferred community structure and extinction selectivity across the Permian–Triassic mass extinction

**DOI:** 10.64898/2026.06.22.733099

**Authors:** Baran Karapunar, Tanya Strydom, Andrew P. Beckerman, William J. Foster, Jennifer A. Dunne, Paul B. Wignall, Pincelli Hull, Catalina Pimiento, Crispin T. S. Little, Andy Ridgwell, Alexander M. Dunhill

## Abstract

The Permian–Triassic mass extinction fundamentally restructured ecosystems, yet it remains unresolved whether ecological collapse unfolded gradually or in discrete steps, and how extinction selectivity varied with environmental change. Here, we analyse marine food webs from Meishan (China) at high temporal resolution. We show that trophic structure destabilised prior to peak biodiversity loss, followed by a structural tipping point during the extinction interval when the community became less robust to secondary extinctions. Extinction selectivity shifted substantially across the study interval. During the extinction interval, extinction was concentrated at lower trophic levels, propagating upwards from benthic herbivores to higher-level consumers. Although coarse resolution preserves temporal trends in trophic structure and extinction selectivity, it obscures abrupt transitions, sequential extinction dynamics, and critical shifts in ecological organisation. We demonstrate that temporal resolution governs inference of extinction dynamics, with important implications for reconstructing past ecological crises and interpreting ecosystem responses to rapid environmental change.

## Introduction

Understanding how ecological communities reorganise over time is central to ecology and evolutionary biology and has become increasingly important in the modern biodiversity crisis, associated with rapidly changing environmental conditions. Temporal evolution of community structure records the interplay between environmental change, species interactions and turnover, offering the opportunity to decipher the processes that govern ecosystem collapse and recovery across timescales. Yet modern observations capture only a small fraction of the history of ecological organisation(*1*). The fossil record preserves biotic responses to climatic and environmental crises far beyond the temporal reach of modern records and provides the only direct empirical basis for evaluating how biodiversity responds to extreme global change over centennial to millennial time scales. The palaeobiological perspective is especially important as contemporary ecosystems confront accelerating anthropogenic warming, deoxygenation, species loss, and habitat disruption that are comparable to those associated with past biodiversity crises(*2*).

Mass extinctions are especially informative natural experiments as they reveal the limits of ecological resilience under rapid environmental stress. Among these events, the Permian–Triassic mass extinction (PTME) was the most severe biotic crisis of the Phanerozoic, eliminating the majority of marine species and fundamentally restructuring Earth’s biosphere(*3*). The extinction coincided with intense environmental perturbation linked to Siberian Traps volcanism, including sea-surface temperature increases of up to 15°C, widespread marine anoxia, ocean acidification, and disruption of primary productivity(*3, 4*). The relative importance and magnitude of these kill mechanisms varied geographically(*5, 6*). Yet despite decades of study, several fundamental questions remain unresolved: how rapidly marine ecosystems destabilised, whether trophic structure collapsed abruptly or stepwise, and how extinction selectivity changed through the crisis interval.

Previous studies suggests that extinction selectivity across the PTME was non-random, structured by physiology, ecology (feeding, motility, tiering, trophic level), morphology, and geographic range(*6–9*). However, most analyses rely on global compilations or coarse temporal resolution that combine taxa from intervals spanning millions of years. Such aggregation can obscure short-lived ecological states, merge distinct extinction pulses, and blur transitions between background, crisis, and recovery dynamics. Although previous studies have identified shifts in selectivity regimes across the Permian– Triassic transition(*7, 10*), they typically compare patterns inferred from pooled assemblages from the latest Permian and from the extinction interval with pooled earliest Triassic assemblages. As a result, current inferences may represent averages of regionally and temporally heterogeneous extinction dynamics.

This challenge is equally severe for reconstructions of trophic structure. Palaeocommunity food-webs built from time-averaged strata have identified substantial ecological reorganization across mass extinctions(*6, 11–14*). Although time-averaging is advantageous for macroecological studies, as it filters out seasonal noise and increases species preservation, it also poses a challenge. Fossil assemblages reconstructed in time-averaged stratigraphic units may include species that did not coexist, potentially creating interactions that never occurred while masking rapid turnover in genuine ecological relationships. Consequently, it remains unclear whether observed shifts in paleocommunity robustness and stability are artifacts of coarse temporal binning or, if genuine, whether they reflect gradual ecological transitions or abrupt threshold behavior.

Resolving these questions requires records with exceptional chronological control. The Meishan section (China) is the Global Stratotype Section and Point for the Permian/Triassic boundary and is among the most intensively studied PTME localities in the world(*15*). High-precision geochronology indicates that the main extinction interval lasted approximately 60 kyr(*16, 17*), while dense stratigraphic sampling documents pronounced faunal turnover through successive beds(*18*). Previous taxonomic analyses support a two-pulse extinction scenario, at latest Changhsingian and at earliest Griesbachian(*19*). The progressive decline in biodiversity across the Permian–Triassic boundary correlates with rising temperatures and reduced primary productivity during the extinction interval(*20*). Meishan therefore provides a unique opportunity to test how ecological structure changed through the crisis at temporal scales relevant to extinction dynamics.

Previous modelling of marine communities from South China, including Meishan, using three coarse temporal bins suggests that food webs collapsed in the earliest Griesbachian and that post-extinction communities were unstable and vulnerable to secondary extinctions(*12*). Reduced robustness to cascading secondary extinctions has also been documented at Meishan, a pattern that appears restricted to tropical communities(*6*). However, because these analyses rely on relatively coarse temporal bins, they imply stepwise rather than abrupt declines in robustness and stability(*6, 12*). Whether marine community collapse involved a discrete tipping point, and the precise timing of such a transition, therefore remains unresolved.

Here, we reconstruct species-level marine trophic networks across the PTME using the most comprehensive species range dataset yet assembled from the Meishan section and the Paleo Food web Inference Model (PFWIM)(*13, 21, 22*). We first reconstructed all potential trophic interaction links using the stratigraphic beds as time bins at the finest temporal resolution, in 71 discrete time bins. Then we simulated 17 trait-informed sequential extinction scenarios based on selectivity against key biological traits. To integrate community structure in selectivity simulations, we incorporated both primary species extinctions and secondary extinctions that were triggered when consumers lost all prey items. We compared these simulations with extinctions in the fossil record to estimate the most plausible primary extinction targets and sequence in each time bin. We repeat these work flow by coarsening the temporal windows (the number of beds in each time bin).

By analysing the community data across multiple temporal resolutions, we evaluate how time averaging influences inference while directly addressing unresolved questions about ecosystem collapse during Earth’s greatest mass extinction. Specifically, we test whether (1) marine trophic networks underwent a discrete breakdown in structure and robustness during the main extinction interval, and, if so, when this ecological tipping point occurred; (2) extinction selectivity shifted between pre-extincton, main extinction, and post-extinction phases, indicating time-varying ecological filtering across the PTME; and (3) coarse temporal aggregation alters inferred food-web structure and trajectories of ecological change, potentially obscuring the timing and nature of ecosystem collapse. Using this approach, we resolve the timing and sequence of ecosystem responses to the PTME at unprecedented temporal resolution, while also demonstrating how resolution itself governs our ability to detect ecological change in the fossil record. Our results highlight the fundamental role of temporal scale in shaping interpretations of community dynamics, with implications not only for reconstructing past crises but also for understanding ecological responses to environmental change in the present.

## Results and discussion

### Changes in community structure and robustness across the PTME

At a high temporal resolution (5–200 kyr bin length), the collapse of marine ecosystems across the Permian–Triassic is a multi-stage process involving early structural destabilisation, then substantial diversity loss followed by a discrete loss of network robustness. Species alpha diversity declined sharply in the late Changhsingian, from ~200–268 species prior to the main extinction interval to 20–40 species afterward (Fig. 1). This biodiversity loss was accompanied by pronounced reorganisation of trophic structure, including increased connectance, mean trophic level, and generality, alongside reduced direct competition.

**Figure 1:**
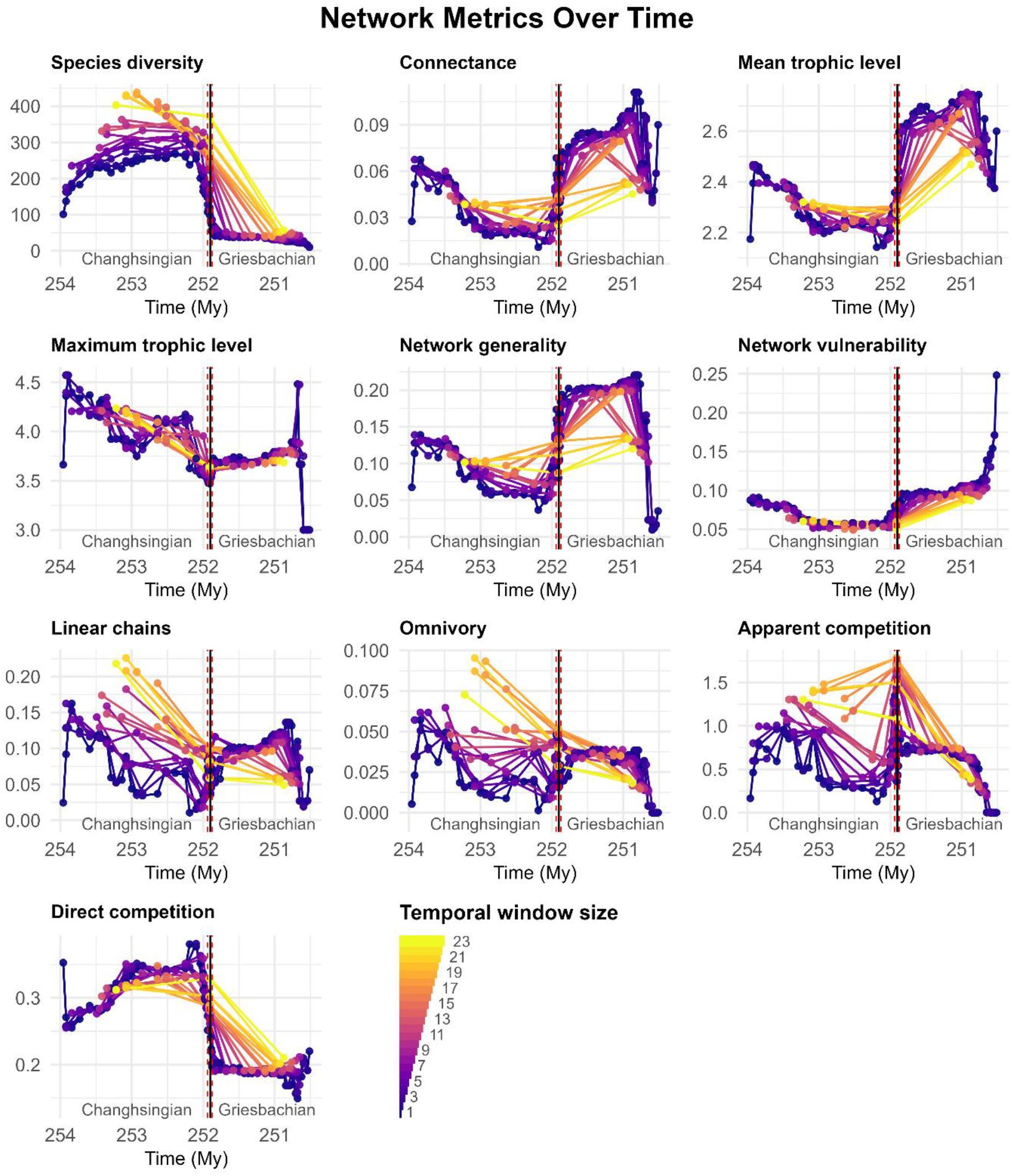
Food web metrics across PTME, measured over different time scales, using equal window sizes (i.e. equal number of beds). Red dashed lines indicate the onset and the end of the main extinction interval (Beds 25–28). Black line indicates the Permian/Triassic boundary.

Interestingly, structural changes preceded peak species loss. Maximum trophic level declined from >4 to <3.5 during the Changhsingian, indicating progressive loss of apex consumers prior to the main extinction interval (between two extinction pulses). This early loss of higher trophic levels suggests that ecosystem destabilisation began before maximum taxonomic loss, implying that trophic structure may have become increasingly fragile leading into the extinction phase rather than collapsing synchronously with it. The early disappearance of top predators from the metacommunities, however, could be partly related to the Signor-Lipps effect(*23*).

Whether this restructuring constitutes true ecosystem collapse depends on how structure or stability is defined. Using maximum trophic level as a proxy, the progressive loss of higher trophic levels implies that structural degradation began in the Changhsingian, contrasting with the interpretation of Huang et al.(*12*) that community collapse occurred in the Early Griesbachian, after the second extinction pulse (Bed 28, top of *Isarcicella staeschei* Zone). However, this metric alone does not capture the system’s susceptibility to cascading extinctions.

A more direct assessment of community resilience to collapse is provided by network robustness, which quantifies vulnerability to secondary extinction cascades. Breakpoint analysis identifies the latest Changhsingian Bed 27b (stage 35) as marking a statistically significant shift in robustness (Fig. 2), indicating a discrete shift in resilience to community collapse. This transition precedes the second extinction pulse, previously defined in Bed 28, and is placed in Bed 27d (stage 37) in our analysis (Table S3, Fig. S4). Hence, the loss of robustness happened during elevated extinction intensity, following a substantial diversity loss, and earlier than previously suggested by coarse temporal analyses (*12*).

**Figure 2:**
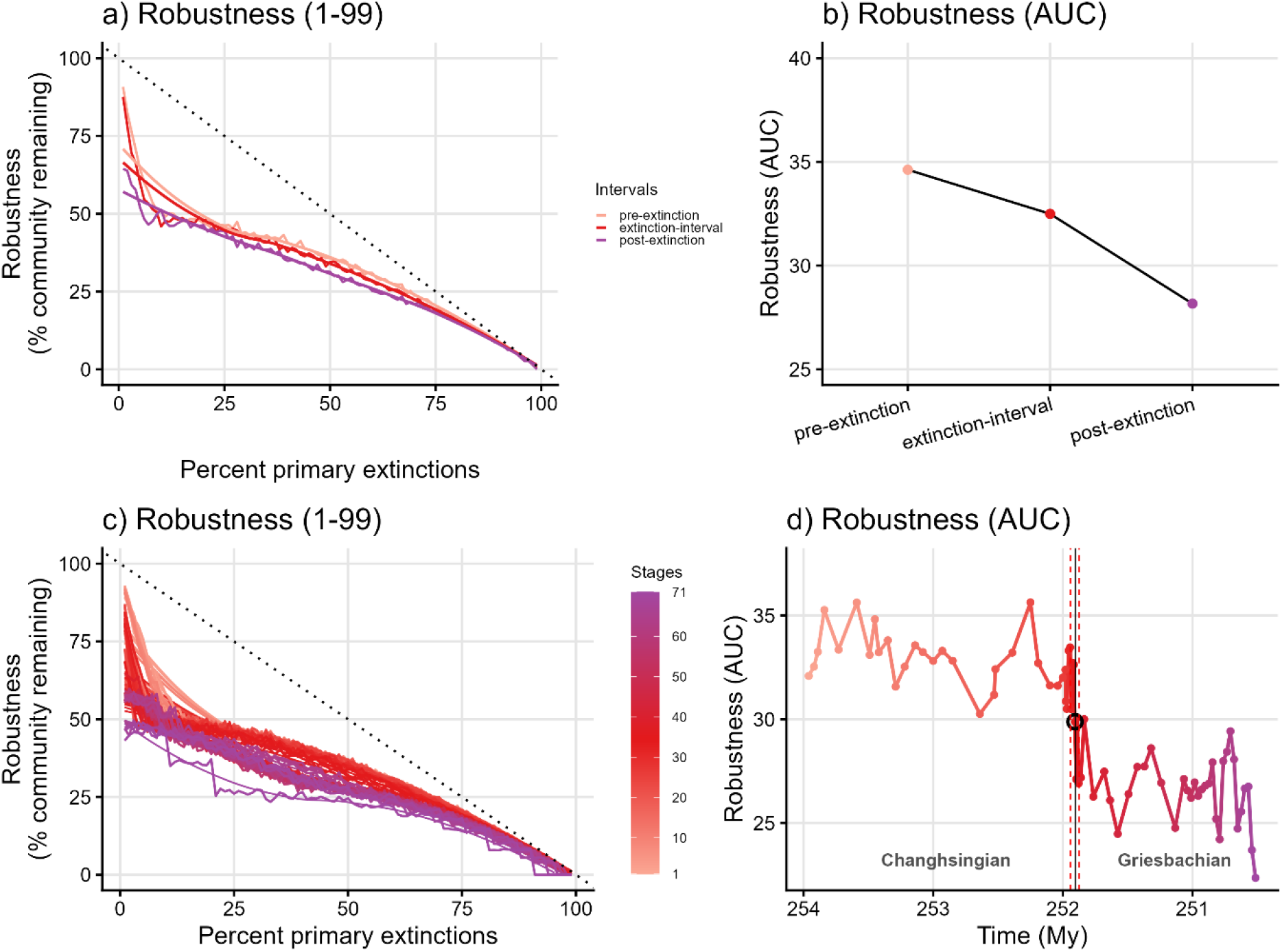
Robustness against random primary extinctions across the PTME. Top panels (a, b) show robustness estimates in low temporal resolution, across three subsequent communities. Bottom panels (c, d) show robustness estimates in high temporal resolution, across 70 subsequent communities. The left panels (a, c) show the Robustness (1–99%) curves drawn by a LOESS fit of mean values after 500 trials. Single data points in the right panels (b, d) represent the area under the curve (AUC) of the Robustness (1–99%) curves. The black circle indicates a statistically significant shift in Robustness in Bed 27c, identified by the breakpoint analysis. Red dashed lines indicate the onset and the end of the main extinction interval. Black line indicates the Permian/Triassic boundary.

The magnitude of change in robustness is modest but significant: robustness declines from approximately 30–36% to 24–30% across Bed 27b. Despite this reduction, post-extinction communities retain few higher trophic nodes (cephalopods and fishes), indicating that trophic networks did not fully collapse but instead transitioned into a persistently less resilient configuration. This suggests that the PTME did not result in complete disintegration of trophic structure, but rather a shift to a degraded yet functionally connected ecosystem.

Temporal resolution affects the visibility of this transition but not its overall trajectory, when window sizes (i.e. number of beds) are roughly equal. Coarse-resolution analyses suggest a gradual and insignificant decline in robustness (Fig. 2b), whereas high-resolution analyses reveal pronounced volatility and a sharp decline concentrated between stages 35–40 (Beds 27b–29b, Fig. 2d). Similarly, early loss of members of higher trophic levels prior to the main extinction phase is only detectable at finer scales (Fig. 1). These comparisons indicate that while broad trends in ecosystem decline are preserved under time averaging, the abruptness and timing of the tipping point are only resolvable at high temporal resolution.

Taken together, these results demonstrate that marine trophic networks did not collapse gradually or solely in the aftermath of extinction pulses. Instead, they underwent early structural destabilisation and biodiversity loss, followed by a discrete breakdown in robustness during the extinction interval, closely aligned with high magnitudes of biodiversity loss. This pattern supports a model in which ecosystem collapse during the PTME involved threshold behaviour, with networks crossing a structural tipping point rather than deteriorating smoothly through time.

### Time-varying extinction selectivity across background, crisis, and recovery phase

Extinction selectivity across the Permian–Triassic transition was dynamic rather than static, shifting between pre-extinction background conditions (Beds 4b–24), the main extinction interval (Beds 25–28), and post-extinction recovery (Beds 29–60). Fine partitioning of the record, rather than comparing only pre- and post-extinction faunas at geological stage level and global scale(*9, 24*), allows direct comparison of these phases, revealing that extinction risk was strongly trait-dependent but varied through time.

During the Changhsingian background interval, extinction selectivity was heterogeneous and temporally unstable (Fig. 3). High-resolution selectivity analyses evaluating selectivity using species identitites (True Skill Statistic) indicate fluctuating targeting across ecological traits, with extinction intermittently affecting both motile and non-motile taxa and often favouring organisms that are poorly buffered against ocean acidification, exhibiting low respiratory capacity and specialist feeding strategies. Mean absolute differences (MAD) between simulated and empirical food web metrics support this pattern, showing shifts from preferential loss of non-motile taxa in the earlier Changhsingian to motile taxa later in the interval. This variability suggests that background extinction operated under weak or shifting ecological filters, rather than a consistent selective regime.

**Figure 3:**
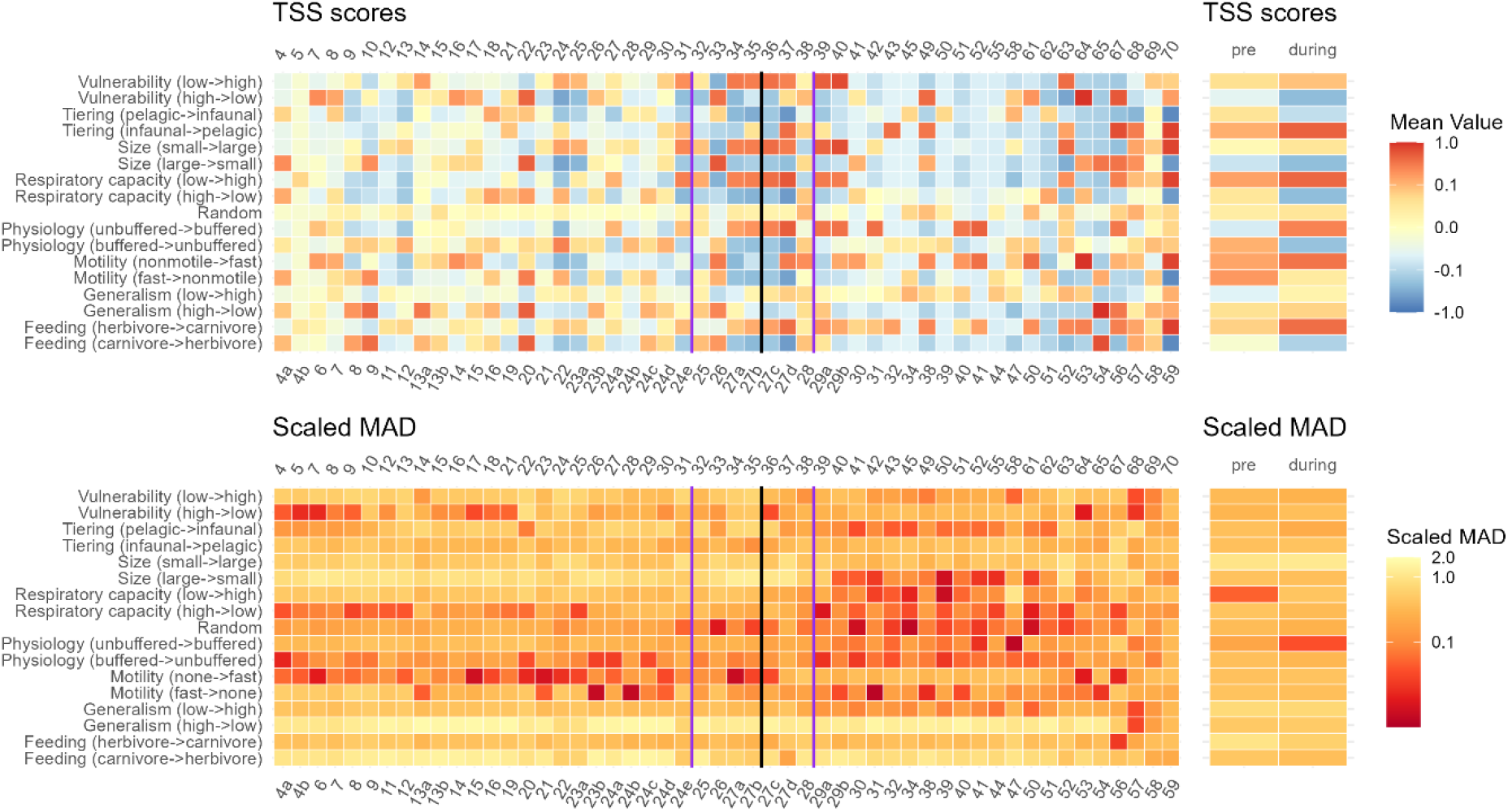
Primary extinction selectivity in high temporal resolution (across 71 time bins) and low resolution (three time bins) using extinction simulations under random scenario and directionally targeting various ecological or physiological traits. The x-axis shows the stages (upper axis) and the corresponding beds (lower axis). Red colour indicates better fit between the simulations and the empirical record, both for the True Skill Statistic (TSS) scores and the scaled mean absolute differences (MAD). Similar colour indicates similar likelihood. Purple lines indicate the onset and the end of the main extinction interval. Black line indicates the Permian/Triassic boundary.

In contrast, the main extinction interval (stages 32–38) is characterised by a more coherent and directional selectivity signal. Both coarse and high-resolution analyses indicate preferential loss of benthic unbuffered herbivores with low respiratory capacity (Fig. 3), consistent with environmental stressors such as warming, deoxygenation and primary productivity decline (Foster et al. 2024). At finer temporal resolution, this signal resolves further: extinction disproportionately affected small-bodied, unbuffered, low-motility taxa, particularly ecological specialists (such as foraminifera and brachiopods). These patterns indicate the emergence of strong physiological filtering during the crisis, replacing the more variable background regime.

Superimposed on this selectivity regime shift is a repeated trophic sequence across both extinction pulses. Initial losses are concentrated among benthic herbivores with low respiratory capacity(in stages 31–32 and 36–37), followed by increased extinction among larger-bodied, higher trophic-level carnivores (in stages 33 and 38, Figure 3). This pattern is consistent with bottom-up propagation of ecological disturbance, in which disruption to primary consumers cascades through the food web to affect higher trophic levels. Both the coherence of the selectivity regime and the repeated sequential selectivity pattern during the main extinction interval suggest an inherent relationship between elevated extinction magnitude and the selectivity regime.

Following the main extinction interval, selectivity shows initial targeting of small-bodied, unbuffered taxa with low vulnerability (foraminifera, bryozoans, ostracods) (in stages 39–40), followed by greater stochasticity in extinction patterns (Figure 3). This indicates a transition to a novel, restructured ecosystem in the post-extinction interval, where ecological interactions and environmental pressures differed from both background and crisis conditions.

Temporal resolution affects how clearly these extinciton selectivity dynamics are resolved but does not fundamentally alter the overarching pattern (Figure 3). Coarse binning captures the broad shift from background variability to crisis-driven physiological filtering, but it compresses distinct ecological phases and obscures within-interval heterogeneity. Only high-resolution analyses reveal the sequential structure of extinction, including fluctuating background selectivity, the emergence of strong crisis filtering, and the repeated bottom-up cascade across extinction pulses.

Together, these results demonstrate that extinction selectivity during the Permian–Triassic transition was non-random and temporally structured, reflecting shifting ecological filters and recovery. Rather than a single extinction regime, the PTME involved multiple, time-varying selective processes. This temporally dynamic extinction selectivity pattern suggests that species survival depends on community structure, species traits, and environmental forcing and that the relative importance of these drivers has changed through time.

Early extinction of top predators prior to the main extinction interval, together with a longer-term decline in the ratio of primary consumers in the Meishan communities (Fig. S3) contrasts somewhat with patterns observed in modern marine ecosystems. In the Adriatic Sea, for example, top predators have declined while primary consumers have increased over the last several millennia as a consequence of human-driven eutrophication and overexploitation(*1*). The decline in diversity at Meishan during the main extinction phase has instead been linked to reduced primary productivity and elevated temperatures(*20*), consistent with our inference that extinction selectivity targeted lower trophic levels and propagated upward through the food web. Despite these contrasting ecological trajectories, both the modern Adriatic(*1*) and the Meishan communities experienced increased vulnerability to secondary extinctions following biodiversity loss. This convergence suggests that ecosystem stability and resilience may be less dependent on which trophic groups are initially lost than on the cumulative erosion of biodiversity and ecological function. Consequently, extinction of different functional groups can drive communities towards similar stability thresholds, increasing the risk of cascading ecological collapse.

### Implications: temporal aggregation and the dynamics of mass extinction

Our results show that time-averaging does not simply reduce resolution, it reshapes our interpretation of extinction dynamics. Across both trophic structure and extinction selectivity, coarse temporal aggregation preserves broad directional trends (e.g. increasing connectance, declining robustness, selectivity toward physiologically constrained taxa) if the dataset is divided into three or more temporal bins, but it systematically obscures the timing, sequence, and mechanisms that underpin ecosystem collapse.

Our inferences are not only influenced by the temporal resolution, but also by our decisions about where to draw bin boundaries. Broad temporal trends between the start and end points are largely preserved in coarse resolution analyses (Figs 1, 4). However, if the previously identified extinction pulses are used to draw bin boundaries between three time bins, the middle time bin representing the main extinction interval becomes temporally very narrow, and this eventually alters the inferred direction of change in some network metrics (connectance, network generality, Fig. 4). Thus, ecological inference and directional trends are sensitive to how temporal boundaries are drawn, even when the number of time bins is kept the same.

**Figure 4:**
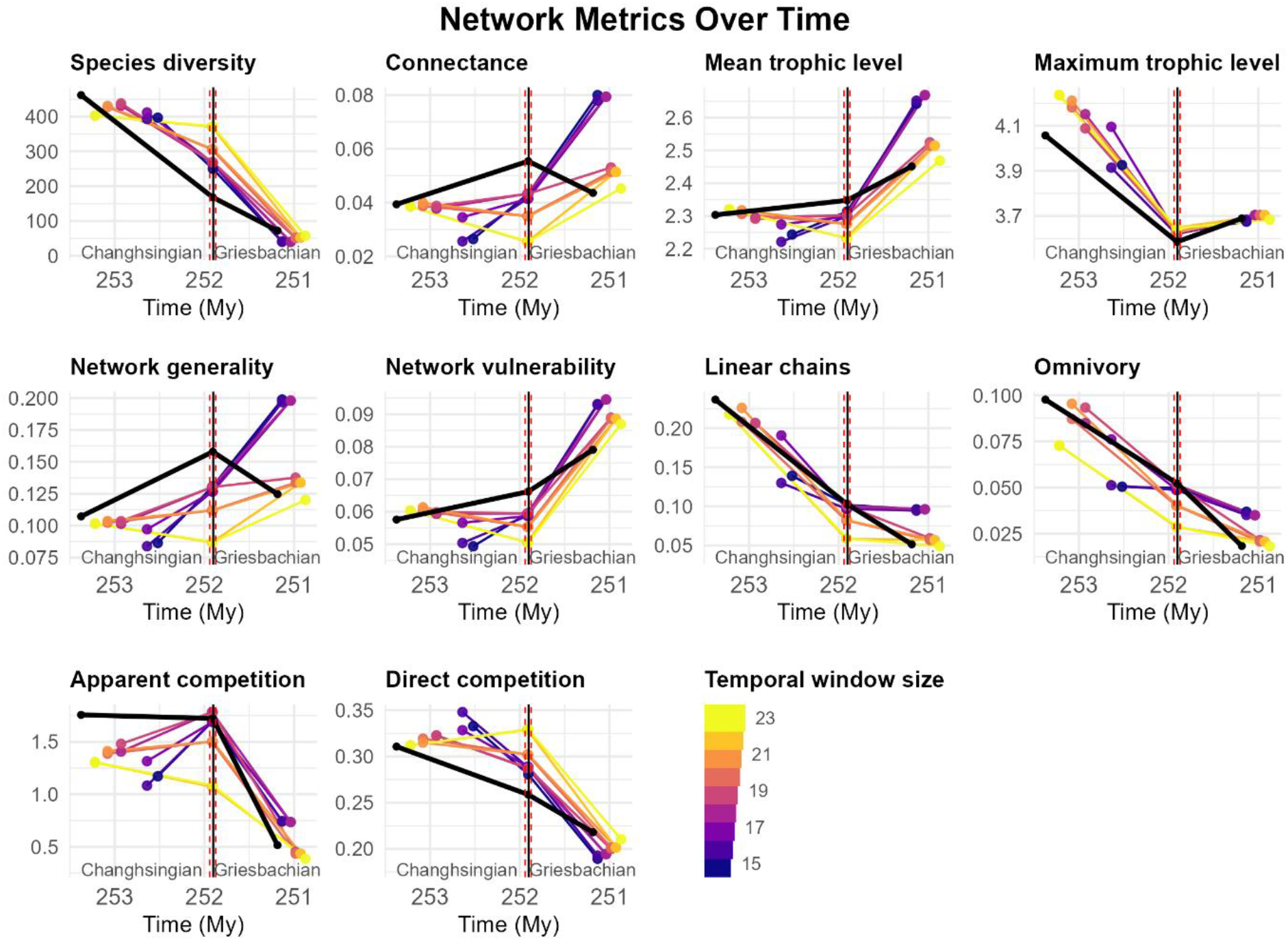
Metrics in three time bins, with equal (colored) and unequal (black) window sizes (number of beds). The unequal binning is based on the previously identified extinction pulses. Red dashed lines indicate the onset and the end of the main extinction interval. Vertical black line indicates the Permian/Triassic boundary.

In trophic networks, time-averaging smooths the transition in robustness that appears threshold-like at high resolution. The discrete breakdown in robustness identified at Bed 27b (stage 35) becomes a gradual decline under coarse binning, masking evidence for tipping-point behaviour. This implies that ecosystems that collapse via thresholds may appear to deteriorate progressively when viewed through temporally aggregated data, leading to underestimation of abrupt ecological change.

Our results show that the selectivity inference depends on the time-scale. Temporal aggregation compresses multiple, temporally distinct selectivity regimes into a single averaged signal. The transition from heterogeneous background selectivity to strongly directional physiological filtering during the extinction interval, to more stochastic post-extinction dynamics is reduced to a simplified contrast between “pre-” and “during-” extinction states. This obscures both the within-interval variability and the sequential, bottom-up propagation of extinction evident at finer scales. While previous studies have emphasized spatial and taxonomic variation in extinction selectivity patterns(*6, 25*), our new results demonstrate that temporal resolution is critical for extinction selectivity inference and fine temporal resolution is essential for resolving the ecological mechanisms underlying extinction dynamics.

Crucially, temporal scale effects are not random loss of information but introduce systematic biases. Time-averaged datasets tend to dampen or intensify ecological volatility (variation in metrics), and blur directional changes and precise timing of community collapse. As a result, key features of mass extinction, such as ecosystem tipping points and temporally shifting selectivity regimes, may be consistently under detected in coarse-resolution records.

More broadly, this suggests that many macroevolutionary inferences derived from temporally aggregated fossil data may be conservative or incomplete, particularly with respect to the pace and structure of ecological collapse. If the PTME is representative of mass extinctions in general, then extinction dynamics are likely to be intrinsically time-structured, involving rapid transitions and shifting ecological filters that cannot be fully resolved without data collected at high stratigraphic resolution. These findings therefore highlight temporal resolution as a critical factor in reconstructing extinction processes, with direct implications for interpreting past crises and caution against using deep-time analogues to understand the ongoing biodiversity loss without incorporating the temporal scale differences. Even our study’s exceptional time resolution of 5–200 kyr is several orders of magnitude longer than the decadal to centennial time scale of current observations(*1*).

## Conclusions

Marine ecosystems across the PTME did not collapse as a single, uniform event, but instead underwent a two-stage transition involving early destabilisation of trophic structure followed by a discrete breakdown in robustness and major biodiversity loss during the extinction interval. The decline in robustness to secondary extinctions indicates that ecosystem collapse was driven not only by reduced species richness, but also by the erosion of key ecological groups that underpin network stability.

Extinction selectivity was strongly time-dependent, shifting between background, crisis, and recovery phases and reflecting changing ecological filters through the course of environmental perturbation. These shifts include repeated patterns of bottom-up disruption propagating through trophic levels, highlighting that extinction was a dynamically structured process rather than a single homogeneous filtering event.

Crucially, these dynamics are only fully resolved at high temporal resolution. While time-averaged data preserve broad trends in diversity and network structure, they systematically obscure abrupt transitions, transient instability, and sequential extinction dynamics. This can lead to an overly gradual representation of what are, in reality, threshold-like ecological changes.

Together, these results show that extinction dynamics are inherently time-structured and mechanistically complex, requiring fine-scale temporal frameworks to resolve the tempo and mode of ecosystem collapse. Temporal resolution therefore plays a central role in shaping interpretations of past extinction events, and caution is warranted when extrapolating from temporally coarse fossil datasets to ecological crises operating on much shorter timescales.

## Supporting information

Supplementary Results

## Author contributions

Conceptualization: BK, TS, APB, and AMD

Data Curation: BK

Formal Analysis: BK, TS

Funding acquisition: APB, JAD, PBW, PH, CP, AR, CTSL, AMD

Investigation: BK

Methodology: BK, TS, APB, JAD, and AMD

Resources: BK

Software: BK, TS, APB, AMD

Supervision: BK, AMD

Validation: BK, TS Visualization: BK

Project administration: BK, APB, AMD

Writing – original draft: BK

Writing – review & editing: BK, TS, APB, WJF, JAD, PBW, PH, CP, AR, CTSL, AMD

## Supplementary Material

Materials and Methods (MM)

Supplementary Results (SR)

Figs. S1 to S5

Tables S1 to S3

References *(MM1*–*MM152)*

