## Supplementary Methods for "Temporal resolution reshapes dynamics of inferred community structure and extinction selectivity across the Permian–Triassic mass extinction"

### Material and Methods

#### Data

The metacommunity datasets were collected from the Meishan Section (Zhejiang, China), which is the most intensely sampled Permian–Triassic section (*1*). The Meishan occurrence and size datasets were compiled from 95 publications (*1-95*). We built our occurrence dataset based on the original dataset used by Foster et al. (*96*), which was originally downloaded from the Geobiodiversity Database (GBDB) and vetted by them. That dataset consists of 603 species. We excluded Wuchiapingian and Dienerian species occurrences and 77 taxa not assigned to species level as well as any occurrence that could not be validated back to an original publication. From the remaining dataset, a total of 152 species were either renamed, or synonymized, or reassigned to another genus based on recent taxonomic knowledge. Finally, the dataset was further improved by adding 102 species from the literature, mostly nektonic taxa, that were missing in the initial compilation. The Meishan dataset compiled by Huang et al. (*72*) includes the fish taxa *Lissodus xiushuiensis* Wang et al., 2007 (*56*), *Polyacrodus jiangxiensis* Wang et al., 2007 (*56*) and cf. *Caturus* in (*55*). There are no published fossils from the Meishan sections (Changxing County in Zhejiang Province) but they have been found in Xiushui County in Jiangxi Province(*55, 56*). Therefore, these taxa were not included in our analyses. We included the actionopterygian *Bobasatrania* from the bed 29a, which was previously erroneously identified as a carbonate fan (*73*), based on the cross section of tooth plate, that is typical of this taxon (*75*). The current compilation exceeded the previous compilations made from Meishan sections ((*24*): 333 species; (*97*): 168 species; (*72*): 172) and comprises 567 Changhsingian–Griesbachian species.

As both the Permian-Triassic boundary and bed numbers differ among the Meishan sections and in different publications, we used the latest framework used by Foster et al.(*96*) which also provides a comprehensive occurrence data of benthic invertebrates from the Meishan sections (Meishan A, B, C, D, E, Z; see (*64*) for more details about the sections). Foster et al.(*96*) updated the stratigraphic information of the occurrences based on latest stratigraphic knowledge (e.g., mixed bed 1 (*42*) represents the Changhsingian following (*64*)) and discarded any occurrences with unreliable stratigraphic information.

We checked all stratigraphic ranges and occurrences of each taxon originally given by Foster et al.(*96*) by referring back to the original references, and then updated these accordingly based on the most recently published records from the Meishan section. To mitigate potential sampling biases, we used species range data and assumed that each taxon was present continuously between its first appearance datum (FAD) and last appearance datum (LAD), even if it was not recorded in every intervening interval. We did not explicitly correct for the Signor–Lipps effect because sampling and preservation potential are unlikely to be uniform across taxa. Confidence interval approaches would tend to overextend the LADs of rare or sparsely sampled groups, such as actinopterygians, which exhibit larger gaps between occurrences, potentially introducing additional bias into extinction timing estimates.

Following Song et al.(*97*), Changhsingian to Griesbachian Beds 25–28 (*changxingensis* and *parvus* conodont Zones) are regarded as the main extinction interval. The Changhsingian Beds from 4b to 24e are regarded as the pre-extinction interval; the Griesbachian Beds 29 to 60 are regarded as the post-extinction interval. As some beds are subdivided by using letters (a.g., 4a, 4b), we assign a number to each of them and named them as "stages". The geochronological data for the Meishan section (Table S1) were taken from Foster et al.(*96*), who provided data up to Bed 50 based on Burgess et al.(*98*) and Shen et al.(*99*). We projected the age assignments from Bed 50 to Bed 60 assuming equal age, as there are no geochronological data available for those beds. Conodont zonation follows Yuan et al.(*100*) and Chen et al.(*12*). The correlating stages of each bed and estimated geochronologic age of each bed are given in Table S1 and in the Supplementary Files and Scripts.

**Table S1.** Generalized stratigraphy and assignments of strata to pre-, during and post-extinction intervals in Meishan, China.

| Period | Age | Formation | Stage | Bed | Conodont zone | Bottom (Myr) | Mid (Myr) | Top (Myr) | Duration (kyr) | Extinction intervals |
|---|---|---|---|---|---|---|---|---|---|---|
| Permian | Wuchiapingian | Longtan | 0 | 1 | *C. longicuspidata* | 254.810 | 254.390 | 253.970 | 84000 | |
| | | | 1 | 1u | | 253.970 | 253.960 | 253.940 | 25000 | |
| | | Changxing | 2 | 2 | | 253.940 | 253.920 | 253.900 | 44167 | |
| | | | 3 | 3 | | 253.900 | 253.890 | 253.880 | 21667 | |
| | | | 4 | 4a | *C. wangi* | 253.880 | 253.840 | 253.800 | 75000 | |
| | Changhsingian | | 5 | 4b | | 253.800 | 253.730 | 253.660 | 141667 | pre-extinction |
| | | | 6 | 5 | | 253.660 | 253.590 | 253.510 | 150000 | |
| | | | 7 | 6 | | 253.510 | 253.490 | 253.470 | 40000 | |
| | | | 8 | 7 | | 253.470 | 253.450 | 253.440 | 27190 | |
| | | | 9 | 8 | | 253.440 | 253.420 | 253.390 | 50030 | |
| | | | 10 | 9 | | 253.390 | 253.350 | 253.310 | 84781 | |
| | | | 11 | 10 | | 253.310 | 253.290 | 253.280 | 27861 | |
| | | | 12 | 11 | *C. subcarinata* | 253.280 | 253.220 | 253.160 | 115938 | |
| | | | 13 | 12 | | 253.160 | 253.140 | 253.110 | 57819 | |
| | | | 14 | 13a | *C. changxingensis* | 253.110 | 253.080 | 253.050 | 54823 | |
| | | | 15 | 13b | | 253.050 | 253.000 | 252.960 | 95866 | |
| | | | 16 | 14 | | 252.960 | 252.930 | 252.910 | 45836 | |
| | | | 17 | 15 | | 252.910 | 252.850 | 252.750 | 164240 | |
| | | | 18 | 16 | | 252.750 | 252.640 | 252.530 | 213987 | |
| | | | 19 | 17 | | 252.530 | 252.530 | 252.530 | 4697 | |
| | | | 20 | 18 | | 252.530 | 252.520 | 252.510 | 20877 | |
| | | | 21 | 19 | | 252.510 | 252.390 | 252.280 | 224426 | |
| | | | 22 | 20 | | 252.280 | 252.250 | 252.220 | 62630 | |
| | | | 23 | 21 | | 252.220 | 252.190 | 252.160 | 61065 | |
| | | | 24 | 22 | | 252.160 | 252.100 | 252.050 | 106153 | |
| | | | 25 | 23a | *C. yini* | 252.050 | 252.040 | 252.020 | 32795 | |
| | | | 26 | 23b | | 252.020 | 252.000 | 251.990 | 32795 | |
| | | | 27 | 24a | | 251.990 | 251.980 | 251.980 | 5045 | |

| | | | | | | | | | | |
|---|---|---|---|---|---|---|---|---|---|---|
| | | | 28 | 24b | | 251.980 | 251.977 | 251.974 | 5550 | |
| | | | 29 | 24c | | 251.974 | 251.970 | 251.966 | 8577 | |
| | | | 30 | 24d | | 251.966 | 251.956 | 251.946 | 19677 | |
| | | | 31 | 24e | *C. meishanensis* | 251.946 | 251.944 | 251.941 | 5045 | |
| | | | 32 | 25 | | 251.941 | 251.938 | 251.934 | 7055 | **extinction** |
| | | | 33 | 26 | *C. zhejiangensis* *H. changxingensis* | 251.934 | 251.927 | 251.920 | 13825 | |
| | | | 34 | 27a | | 251.920 | 251.916 | 251.911 | 9217 | |
| | | | 35 | 27b | | 251.911 | 251.906 | 251.902 | 9217 | |
| **Triassic** | **Griesbachian** | **Yinkeng** | 36 | 27c | *H. parvus* | 251.902 | 251.897 | 251.893 | 9217 | |
| | | | 37 | 27d | | 251.893 | 251.888 | 251.883 | 9217 | |
| | | | 38 | 28 | *I. staeschei* | 251.883 | 251.879 | 251.874 | 9663 | |
| | | | 39 | 29a | | 251.874 | 251.861 | 251.848 | 25575 | **post-extinciton** |
| | | | 40 | 29b | *I. isarcica* | 251.848 | 251.835 | 251.823 | 25575 | |
| | | | 41 | 30 | *C. planata* | 251.823 | 251.765 | 251.708 | 115090 | |
| | | | 42 | 31 | | 251.708 | 251.681 | 251.654 | 53708 | |
| | | | 43 | 32 | | 251.654 | 251.636 | 251.618 | 35806 | |
| | | | 44 | 33 | | 251.618 | 251.577 | 251.535 | 82642 | |
| | | | 45 | 34 | | 251.535 | 251.494 | 251.452 | 83017 | |
| | | | 46 | 35 | | 251.452 | 251.426 | 251.399 | 53156 | |
| | | | 47 | 36 | | 251.399 | 251.371 | 251.343 | 56478 | |
| | | | 48 | 37 | | 251.343 | 251.319 | 251.296 | 46512 | |
| | | | 49 | 38 | | 251.296 | 251.241 | 251.187 | 109635 | |
| | | | 50 | 39 | | 251.187 | 251.133 | 251.080 | 106312 | |
| | | | 51 | 40 | | 251.080 | 251.066 | 251.053 | 27628 | |
| | | | 52 | 41 | | 251.053 | 251.039 | 251.025 | 27628 | |
| | | | 53 | 42 | | 251.025 | 251.011 | 250.997 | 27628 | |
| | | | 54 | 43 | | 250.997 | 250.984 | 250.970 | 27628 | |
| | | | 55 | 44 | | 250.970 | 250.956 | 250.942 | 27628 | |
| | | | 56 | 45 | | 250.942 | 250.928 | 250.914 | 27628 | |
| | | | 57 | 46 | | 250.914 | 250.901 | 250.887 | 27628 | |
| | | | 58 | 47 | | 250.887 | 250.873 | 250.859 | 27628 | |
| | | | 59 | 48 | | 250.859 | 250.845 | 250.832 | 27628 | |
| | | | 60 | 49 | | 250.832 | 250.818 | 250.804 | 27628 | |
| | | | 61 | 50 | | 250.804 | 250.790 | 250.776 | 27628 | |
| | | | 62 | 51 | | 250.776 | 250.763 | 250.749 | 27628 | |
| | | | 63 | 52 | | 250.749 | 250.735 | 250.721 | 27628 | |
| | | | 64 | 53 | | 250.721 | 250.707 | 250.693 | 27628 | |
| | | | 65 | 54 | | 250.693 | 250.680 | 250.666 | 27628 | |
| | | | 66 | 55 | *Nc. discreta* | 250.666 | 250.652 | 250.638 | 27628 | |
| | | | 67 | 56 | | 250.638 | 250.624 | 250.611 | 27628 | |
| | | | 68 | 57 | | 250.611 | 250.597 | 250.583 | 27628 | |
| | | | 69 | 58 | | 250.583 | 250.569 | 250.555 | 27628 | |
| | | | 70 | 59 | | 250.555 | 250.541 | 250.528 | 27628 | |
| | | | 71 | 60 | | 250.528 | 250.514 | 250.500 | 27628 | |

According to PALEOMAP plate model (*101*) using the chronosphere R package (*102, 103*), the Meishan section was located at a palaeolatitude of 17°N. The Meishan deposits are interpreted to have been deposited on slope of a carbonate platform (Zhang et al. in (*1, 97, 104*)), with the environment becaming shallower towards the main extinction horizon (subtidal, below fair-weather wave base) before deepening into the post-extinction interval (Zhang et al. in (*1, 12, 104*)). Beds 23 and 24 have been interpreted to contain allochthonous deposits (*12, 105*); however, it is difficult to verify from the literature whether any of the occurrences were transported from nearby environments. Herein, all sampled occurrences within each bed are considered to be part of the metacommunity of that bed.

#### Ecological traits

Each species was assigned four ecological traits which are used to parameterise the food web model: tiering, motility, feeding, following Bambach's Ecospace model(*106*), and a discrete body-size trait. We simplified categories when finer distinctions did not affect trophic interpretations: surficial and erect forms were grouped as epifaunal; cemented, byssate, and other attached taxa were treated as "attached"; deposit-, filter-, and grazer-herbivores were combined as "herbivores"; and both grazer-carnivores (such as pleurotomariid gastropods) and active hunters were all assigned to a single "carnivore" category. Keeping a single carnivore category is largely because direct evidence of dietary specialism is challenging to infer from the fossil record, as fossilized gut contents for specific taxa are extremely rare, and more commonly fossilized coprolites are difficult to attribute to specific producers at lower taxonomic levels. Furthermore, available gut contents and coprolites of fossil predators suggest diverse diets within the same clade, as seen in both Vertebrata(*107-111*) and Cephalopoda(*112*). Our approach may overestimate the potential prey links of carnivore taxa by assuming generalism. However, this approach also avoids the artificial exclusion of carnivorous taxa from the reconstructed food webs. Imposing overly restrictive dietary specialisation can result in some carnivores being assigned no viable prey species, causing them to be disconnected from the network and subsequently excluded from the trophic web.

Ecological traits were interpreted based on functional morphology, phylogenetic affinity, direct evidence (such as gut contents, isotopes), and predation traces (such as drill holes, scratch or bite marks) according to the published literature ((*106, 113-115*), Bivalvia(*116, 117*), Gastropoda(*118-122*), Cephalopoda(*112, 123, 124*), Echinodermata(*125-127*), Brachiopoda(*128-130*), Ostracoda(*131*), Bryozoa(*132*), Microconchida(*133*), Foraminifera(*134*), Conodonta(*135-137*)).

Size data were collected from the literature, either from the measurements given in the publication or by measuring from the published drawings/photographs or by digitizing individual datapoints from scatter plots. We used linear measurements, i.e. height and length, as proxies for size as they generally represent the primary axes of body size. In some groups (e.g. those with radial symmetry i.e. Ophiuroidea and Anthozoa; as well as some ichnofossils), a single measurement is used as both height and length. Body size categories were assigned using species mean size, calculated as $\sqrt{(\text{height} \times \text{length})}$

for each specimen. Species were classified as tiny (<1 mm), small (1–10 mm), medium (10–100 mm), large (100–1000 mm), or huge (>1000 mm). Shell size was used as a proxy for body size in molluscs, arthropods, foraminifera, and ostracods, burrow diameter for ichnofossils, and individual polyp or zooid size for colonial taxa (bryozoans, corals). For articulated groups (e.g., vertebrates and echinoderms), size estimates were based on disarticulated elements (e.g., conodonts, teeth) following the approach described by Karapunar et al.(*103*).

In addition to the ecological traits used to model trophic interaction, we also compiled the physiological traits of respiratory protein and physiological buffering capability (against decreasing carbonate ions [$CO_3^{2-}$]) to inform primary extinction selectivity in the secondary extinction cascade models. Respiratory protein data for each taxonomic group was gathered from the literature(*138-141*). Physiological buffering of each taxonomic group was assigned according to the division given by Bambach et al.(*142*) and Knoll et al.(*143*). The assigned ecological traits of each species are given in the Supplementary Files and Scripts.

#### Food webs and food web metrics

We used the Paleo Food web Inference Model (PFWIM) (*144-147*) to reconstruct marine metacommunities. Food webs were modelled based on four ecological trait categories (i.e. feeding, motility, tiering, body size) and feeding rules that determine feeding links between organisms (Figure S1). PFWIM reconstructs all potential interactions for all communities in each time bin. The feeding rules were decided by considering modern interactions and ecological theory, which indicates allometric scaling between predator and prey size(*148*). Our discrete coding allows carnivores (representing generalists, large prey specialists and small prey specialist sensu García-Oliva & Wirtz(*149*)) to feed on similar- to smaller-sized individuals, but also on larger-sized prey within their own category. This allows for rare potential cases, where carnivores feed on similar-sized or larger prey taxa. Taxa that feed on prey smaller than 1 mm are coded as microcarnivores (preying on zooplankton), which includes planktivorous taxa that do not follow the allometric size rule(*149*).

We reconstructed feasible metawebs, representing all biologically plausible feeding interactions, for each metacommunity within a time bin. This approach allows for potential trophic rewiring following prey loss, and therefore provides a conservative baseline for modelling extinction cascades, in which species can potentially shift diet within their realised interaction set as prey become extinct. We prefer this approach over realised web reconstruction methods, which generate food webs by stochastic downsampling of feasible interactions to match observed link distributions. While these realised webs aim to reflect empirical network structure, the randomisation of link assignment introduces variability among reconstructed webs, even within the same time bin. This complicates statistical comparison of extinction simulations through time, as differences in cascade dynamics may reflect stochastic realisations rather than ecological signal. In contrast, feasible metawebs provide a consistent and maximally connected baseline, against which differences in extinction vulnerability among communities and time intervals can be more robustly compared(*144*). Although this approach may increase network connectivity and therefore yield more conservative estimates of secondary extinction risk, it reduces artefactual variation and avoids underestimating the capacity of communities for trophic reorganisation following species loss. Finally, we removed cannibalistic interactions from the reconstructed webs, as these can generate unrealistic trophic metrics and artificially depress estimated trophic levels of affected taxa.

The food webs are reconstructed at the species-level, and we added one primary producer node and a zooplankton node. The zooplankton node incorporates larval stage of many marine taxa in the ecosystem, which are feeding on phytoplankton and other zooplankton. All primary consumers (including the zooplankton) have a single feeding link (edge) with the primary producer node.

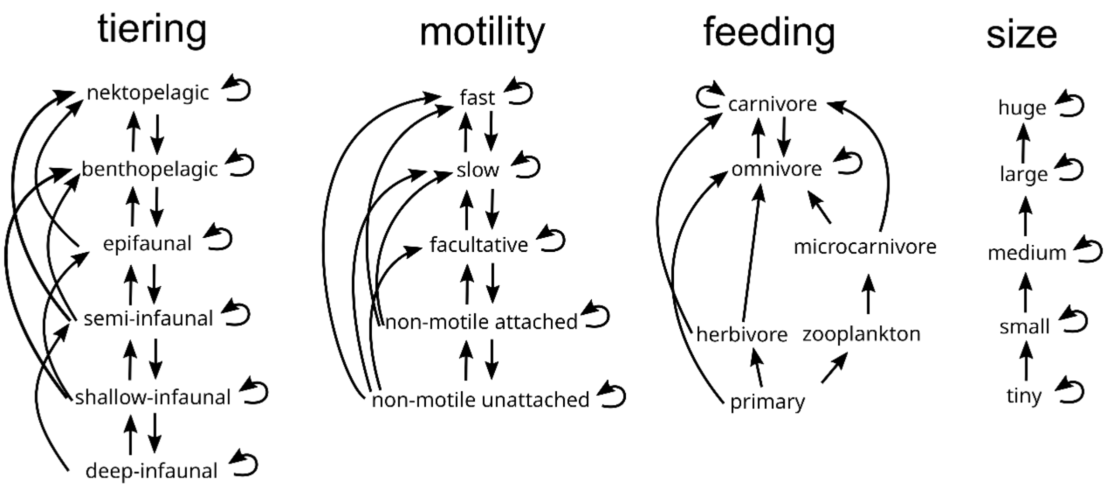


**Figure S1.** Feeding rules to reconstruct food webs using PFWIM.

Food webs were reconstructed first for each of the 71 stages, hence at temporal window size at one. Then, food webs were reconstructed by temporally aggregating the data in discrete non-overlapping windows. The temporal window size was increased gradually from a single stage (71 time bins) to a maximum of 23 stages (3 time bins). The food webs were reconstructed for species that are aggregated in these time windows. When increasing the window size, the stage 35 was taken as a central point, and window sizes were expanded symmetrically. For instance when the window size was 2, the divisions were as (...33+34, 35+36, 37+38...); and when the window size was three, the divisions were (...30+31+32, 33+34+35, 36+37+38...). Then food webs were reconstructed for three consecutive communities, considering the former division of mass extinciton interval(*97*), by assigning species from stages 5–31 (=Beds 4b–24d) to the pre-extinction interval community; species from stages 32–38 (=Beds 25–28) to the main extinction interval community; species from stages 39–71 (=Beds 29 to 60) to the post-extinction community. A total of 246 food webs were reconstructed at various temporal window sizes. The R scripts used for the PFWIM can be found in the Supplementary Files and Scripts. Species compositions in each time bin and the trophic levels estimated by PFWIM are given as html files in the Supplementary Files and Scripts.

**Table S2.** Food web metrics inferred using the PFWIM.

| Metric type | Metric | Description |
|---|---|---|
| Network structure | Connectance | Proportion of possible links in a network that actually occur. Calculated as the number of links (L) divided by number of species (S) squared (i.e. L/S^2). |
| | Species diversity | Number of nodes (species) in a network |
| | Network generality | Standard deviation of normalized in-degree of nodes (see node position metrics below) |
| | Network vulnerability | Standard deviation of normalized out-degree of nodes (see node position metrics below) |
| | Mean trophic level | Mean trophic level of nodes (see node position metrics below) |
| | Maximum trophic level | Maximum trophic level of nodes (see node position metrics below) |
| | Motif: Linear chains | Number of motifs (3-node subnetworks) describing simple linear chains, normalized to S^2. |
| | Motif: Omnivory | Number of motifs describing predators preying on two taxa at different trophic levels, normalized to S^2. |
| | Motif: Apparent competition | Competition between two prey with a common predator. Calculated as the number of motifs describing predators preying on two species, normalized to S^2. |
| | Motif: Direct competition | Competition between two predators with a common prey. Number of motifs describing two predators sharing a prey species, normalized to S^2. |
| Node position | Generality (normalized in-degree) | How many resource nodes a consumer node has. Normalized to the mean number of links per node for that network. |
| | Vulnerability (normalized out-degree) | How many consumer nodes a resource node has. Normalized to the mean number of links per node for that network. |
| | Trophic level | 1 + the weighted mean of the trophic levels of its resources. Primary producers are assumed to have trophic level of 1. |

#### Robustness simulations

Robustness ($R_x$) was calculated for each network using the framework introduced in Jonsson et al.(*150*) where robustness is defined as the proportion of primary extinctions that will result in x% of all species in the network going extinct (from both primary and secondary cascading extinctions). Here, we estimate $R_x$ for x in 1% to 99% in steps of 1%. This was repeated 500 times using a random primary extinction sequence, this was then used to calculate the mean robustness at each x% value.

A low value of $R_x$ for a particular sequence (relative to other sequences) implies a lower robustness as only a small proportion of primary extinction will lead to an x% collapse of the trophic network. An $R_x$ value of x/100 indicates that no secondary extinctions have occurred and 1/S shows that only one primary species deletion is needed to cause an x% collapse of the network.

The robustness simulations were first done for three consecutive communities, pre-extinction (species occurring in stages 5–31), extinction (32–38), and post-extinction (stages 39–71). Then the robustness simulations were conducted for each single stage of 71 consecutive stages. Then we calculated the area under the Robustness (1–99%) curves. We then applied a breakpoint analysis on Robustness (AUC) values across 71 consecutive stages using the "strucchange" R package, which uses an algorithm to find breakpoints in linear regression lines in a time series(*151*). Robustness simulation scripts written in the Julia language, and the breakpoint analysis script in R can be found in the Supplementary Files and Scripts.

#### Extinction selectivity analysis

Extinction cascades were simulated by subjecting species to primary extinction scenarios based on ecological and trophic traits that correspond to known sensitivities to environmental changes (e.g. high temperatures, deoxygenetion) that have been hypothesised as plausible drivers of mass extinction across the PTME. Specifically, we explored 13 different extinction scenarios. We have simulated the primary extinction order under the following scenarios: 1) random, 2) body size (large to small/small to large), 3) tiering (infaunal to pelagic/pelagic to infaunal), 4) motility (fast to non-motile/non-motile to fast), 5) physiology (buffered to unbuffered/unbuffered to buffered [against decreasing carbonate ions]), 6) feeding (carnivorous to herbivorous/herbivorous to carnivorous), 7) respiratory capacity (low to high/high to low), 8) generality (low to high/high to low), and 9) vulnerability (low to high/high to low). Generality and vulnerability of taxa were estimated based on the food webs reconstructed with the PFWIM.

For each replicate, we catalogued the primary extinction and any secondary extinctions that arose. Secondary extinctions were treated as being cascading, whereby after the removal of (a) species, all species that had lost all their prey were deemed to have gone extinct and were also removed. This process was repeated until no more species could be removed. Extinctions were stopped when the diversity of the simulated post-extinction community reached the richness that equalled or less than that of the empirical post-extinction community. We generated 50 replicates for each scenario by sampling randomly among species from within each traits' levels in the sequence. Simulated post-extinction food webs were then compared to the empirical post-extinction community using the following approaches.

For evaluating selectivity scenarios, simulated surviving communities were compared with an empirical surviving community, that is the set of species that are known as upper boundary crossers. The selectivity analysis conducted for two boundaries (between pre/during, and during/post), and then for stage boundaries between 71 stages which recorded extinctions. To test statistical fitting of simulated primary extinction scenarios to the empirical fossil record, we used node identities and trophic metric values.

First, we compared four structural metrics (Connectance, Network generality, Network vulnerability, Maximum trophic level) as well as the frequency of four motifs (Linear chains, Omnivory, Apparent competition, Direct competition) between the empirical and simulated networks (Table S2). This was done by using the mean absolute difference (MAD) between the empirical and simulated networks. We then applied z-scoring to mean absolute differences to scale metric values, in order to prevent a single metric from being weighted disproportionately. A value of zero would indicate that the simulated network perfectly matches the network metrics of the empirical network.

Additionally, we also calculated the True Skill Statistic(*152*) to compare the node level similarities of identity between the empirical and simulated networks:

$$\mathrm{TSS} = \frac{\mathrm{TP} * \mathrm{TN} - \mathrm{FP} * \mathrm{FN}}{(\mathrm{TP} + \mathrm{FN}) * (\mathrm{FP} + \mathrm{TN})}$$

True Positives (TP): Correctly predicted presences (simulated survivors in empirical survivors)
True Negatives (TN): Correctly predicted absence (simulated extinctions in empirical extinction)
False Positives (FP): Incorrectly predicted presences (empirical extinctions in simulated survivors)
False Negatives (FN): Incorrectly predicted absences (simulated extinctions in empirical survivors)
The TSS can range between -1 and 1, where one indicates a perfect match between empirical and simulated network compositions, and anything below zero indicates a performance no better than random.

TSS calculations consider only the identity of species (i.e., has the extinction scenario removed/retained the species known to survive into empirical post-extinction community). We calculated the number of extinct species as the difference between the number of boundary crossers (survivors) and the pre-extinction diversity. We then used this value as the simulated extinction threshold and compared the simulated survivors with the empirical survivors – this means that the TSS calculations will only reflect the removal or retainment of the 'wrong species'. If there is a complete extinction, TSS scores cannot be calculated. We combined the inference from these comparisons to identify the most plausible set of extinction scenarios that could deliver a community that most closely resembles the post-extinction community. The R scripts used for the extinction selectivity analyses can be found in the Supplementary Files and Scripts.
