## Supplementary Results for "Temporal resolution reshapes dynamics of inferred community structure and extinction selectivity across the Permian–Triassic mass extinction"

**Table S3:** Number of extinctions, originations, upper boundary crossers, and the extinction magnitude in each time bin.

| Stage | Bed | Age (Myr) | Extinctions | Boundary crossers | Originations | Extinction magnitude |
|---|---|---|---|---|---|---|
| 1 | 1u | 253.96 | 0 | 99 | NA | 0.00 |
| 2 | 2 | 253.92 | 0 | 135 | 36 | 0.00 |
| 3 | 3 | 253.89 | 0 | 161 | 26 | 0.00 |
| 4 | 4a | 253.84 | 3 | 170 | 12 | 1.73 |
| 5 | 4b | 253.73 | 1 | 180 | 11 | 0.55 |
| 6 | 5 | 253.59 | 0 | 210 | 30 | 0.00 |
| 7 | 6 | 253.49 | 6 | 223 | 19 | 2.62 |
| 8 | 7 | 253.45 | 4 | 224 | 5 | 1.75 |
| 9 | 8 | 253.42 | 17 | 217 | 10 | 7.26 |
| 10 | 9 | 253.35 | 19 | 208 | 10 | 8.37 |
| 11 | 10 | 253.29 | 0 | 214 | 6 | 0.00 |
| 12 | 11 | 253.22 | 8 | 225 | 19 | 3.43 |
| 13 | 12 | 253.14 | 24 | 201 | 0 | 10.67 |
| 14 | 13a | 253.08 | 7 | 235 | 41 | 2.89 |
| 15 | 13b | 253 | 8 | 232 | 5 | 3.33 |
| 16 | 14 | 252.93 | 6 | 240 | 14 | 2.44 |
| 17 | 15 | 252.85 | 12 | 240 | 12 | 4.76 |
| 18 | 16 | 252.64 | 8 | 248 | 16 | 3.13 |
| 19 | 17 | 252.53 | 0 | 254 | 6 | 0.00 |
| 20 | 18 | 252.52 | 0 | 254 | 0 | 0.00 |
| 21 | 19 | 252.39 | 13 | 253 | 12 | 4.89 |
| 22 | 20 | 252.25 | 29 | 229 | 5 | 11.24 |
| 23 | 21 | 252.19 | 11 | 225 | 7 | 4.66 |
| 24 | 22 | 252.1 | 65 | 188 | 28 | 25.69 |
| 25 | 23a | 252.04 | 37 | 162 | 11 | 18.59 |
| 26 | 23b | 252 | 2 | 161 | 1 | 1.23 |
| 27 | 24a | 251.98 | 17 | 156 | 12 | 9.83 |
| 28 | 24b | 251.9772 | 5 | 152 | 1 | 3.18 |
| 29 | 24c | 251.9702 | 4 | 150 | 2 | 2.60 |
| 30 | 24d | 251.956 | 11 | 144 | 5 | 7.10 |
| 31 | 24e | 251.9437 | 56 | 90 | 2 | 38.36 |
| 32 | 25 | 251.9376 | 13 | 92 | 15 | 12.38 |
| 33 | 26 | 251.9272 | 26 | 87 | 21 | 23.01 |
| 34 | 27a | 251.9157 | 18 | 91 | 22 | 16.51 |
| 35 | 27b | 251.9064 | 20 | 76 | 5 | 20.83 |
| 36 | 27c | 251.8972 | 9 | 70 | 3 | 11.39 |
| 37 | 27d | 251.888 | 28 | 44 | 2 | 38.89 |
| 38 | 28 | 251.8786 | 9 | 42 | 7 | 17.65 |
| 39 | 29a | 251.861 | 8 | 39 | 5 | 17.02 |
| 40 | 29b | 251.8354 | 2 | 37 | 0 | 5.13 |

| | | | | | | |
|---|---|---|---|---|---|---|
| 41 | 30 | 251.765 | 1 | 37 | 1 | 2.63 |
| 42 | 31 | 251.6806 | 2 | 36 | 1 | 5.26 |
| 43 | 32 | 251.6359 | 1 | 35 | 0 | 2.78 |
| 44 | 33 | 251.5767 | 0 | 35 | 0 | 0.00 |
| 45 | 34 | 251.4938 | 1 | 34 | 0 | 2.86 |
| 46 | 35 | 251.4257 | 0 | 35 | 1 | 0.00 |
| 47 | 36 | 251.3709 | 0 | 36 | 1 | 0.00 |
| 48 | 37 | 251.3194 | 0 | 36 | 0 | 0.00 |
| 49 | 38 | 251.2414 | 3 | 33 | 0 | 8.33 |
| 50 | 39 | 251.1334 | 1 | 32 | 0 | 3.03 |
| 51 | 40 | 251.0664 | 2 | 30 | 0 | 6.25 |
| 52 | 41 | 251.0388 | 1 | 29 | 0 | 3.33 |
| 53 | 42 | 251.0112 | 0 | 29 | 0 | 0.00 |
| 54 | 43 | 250.9835 | 0 | 29 | 0 | 0.00 |
| 55 | 44 | 250.9559 | 1 | 28 | 0 | 3.45 |
| 56 | 45 | 250.9283 | 0 | 28 | 0 | 0.00 |
| 57 | 46 | 250.9007 | 0 | 28 | 0 | 0.00 |
| 58 | 47 | 250.873 | 5 | 24 | 1 | 17.24 |
| 59 | 48 | 250.8454 | 0 | 24 | 0 | 0.00 |
| 60 | 49 | 250.8178 | 0 | 24 | 0 | 0.00 |
| 61 | 50 | 250.7901 | 1 | 23 | 0 | 4.17 |
| 62 | 51 | 250.7625 | 2 | 22 | 1 | 8.33 |
| 63 | 52 | 250.7349 | 10 | 22 | 10 | 31.25 |
| 64 | 53 | 250.7073 | 2 | 21 | 1 | 8.70 |
| 65 | 54 | 250.6796 | 2 | 19 | 0 | 9.52 |
| 66 | 55 | 250.652 | 0 | 20 | 1 | 0.00 |
| 67 | 56 | 250.6244 | 3 | 17 | 0 | 15.00 |
| 68 | 57 | 250.5967 | 4 | 17 | 4 | 19.05 |
| 69 | 58 | 250.5691 | 5 | 13 | 1 | 27.78 |
| 70 | 59 | 250.5415 | 6 | 8 | 1 | 42.86 |
| 71 | 60 | 250.5138 | NA | NA | 0 | NA |

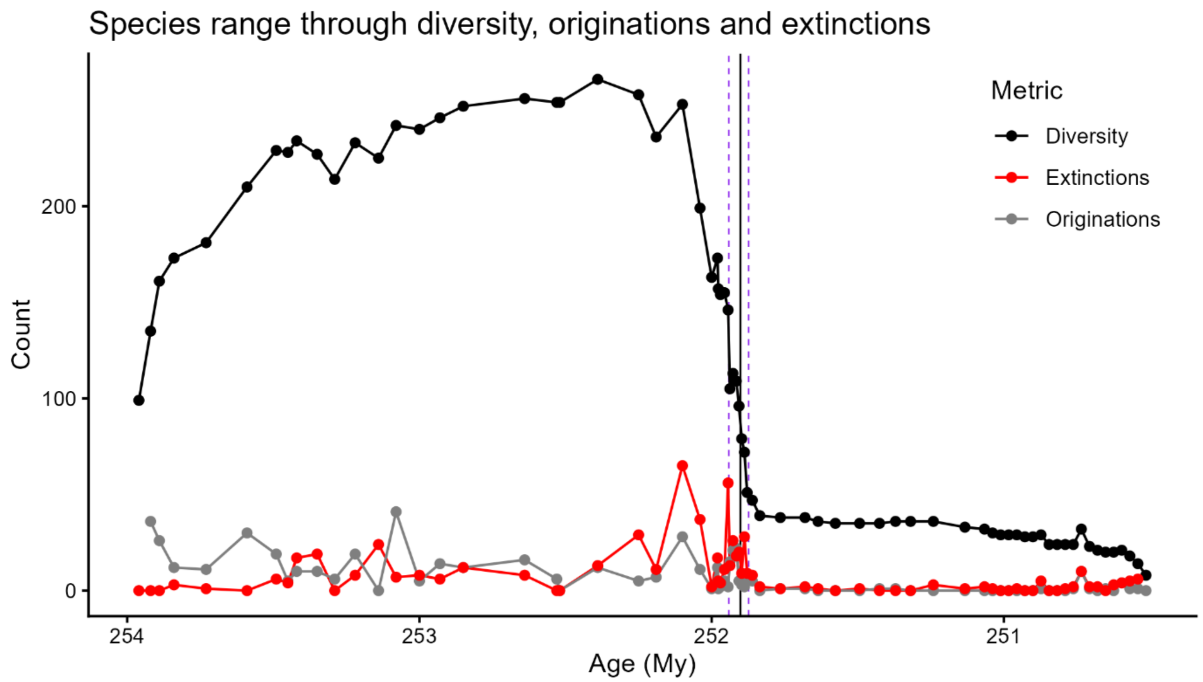


**Figure S2:** Species diversity, number of originations (count of first appearance dates) and extinctions (count of last appearance dates) in each time bin. Purple dashed lines indicate the onset and end of the extinction interval. Black vertical line indicates the Permian–Triassic boundary.

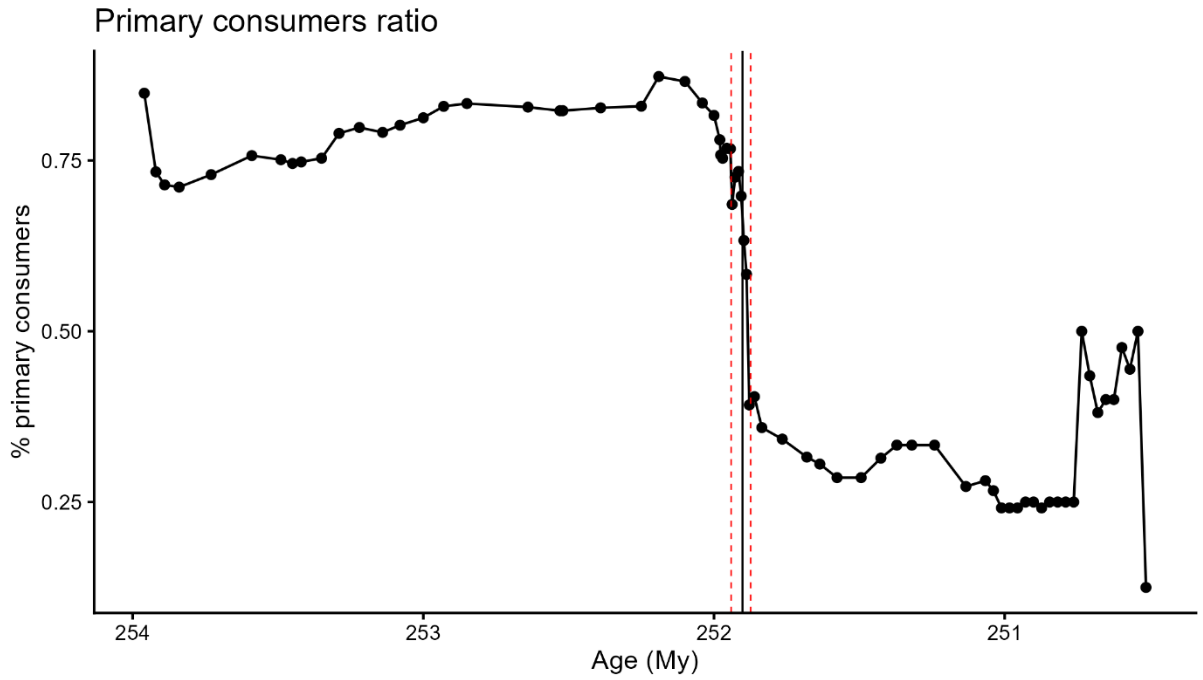


**Figure S3:** Primary consumers ratio (the percentage of herbivores to all other groups) in each time bin. Red dashed lines indicate the onset and end of the extinction interval. Black vertical line indicates the Permian–Triassic boundary.

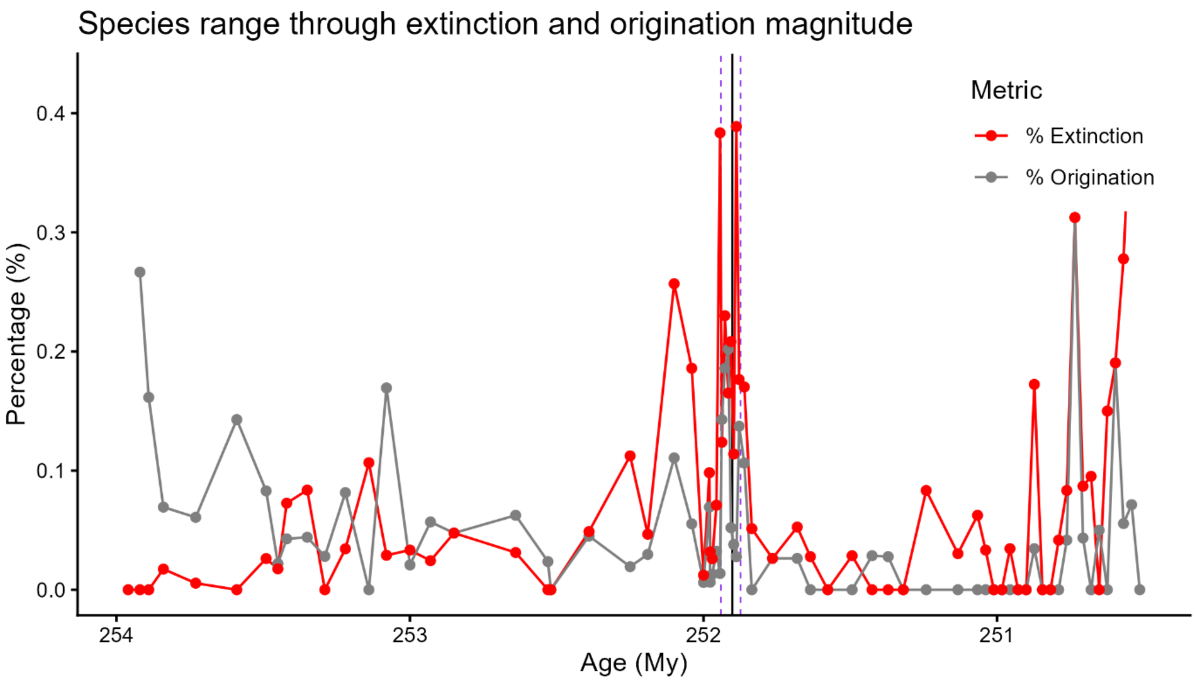


**Figure S4:** Percentage of originations and extinctions relative to standing diversity per time bin. Purple dashed lines indicate the onset and end of the extinction interval. Black vertical line indicates the Permian–Triassic boundary.

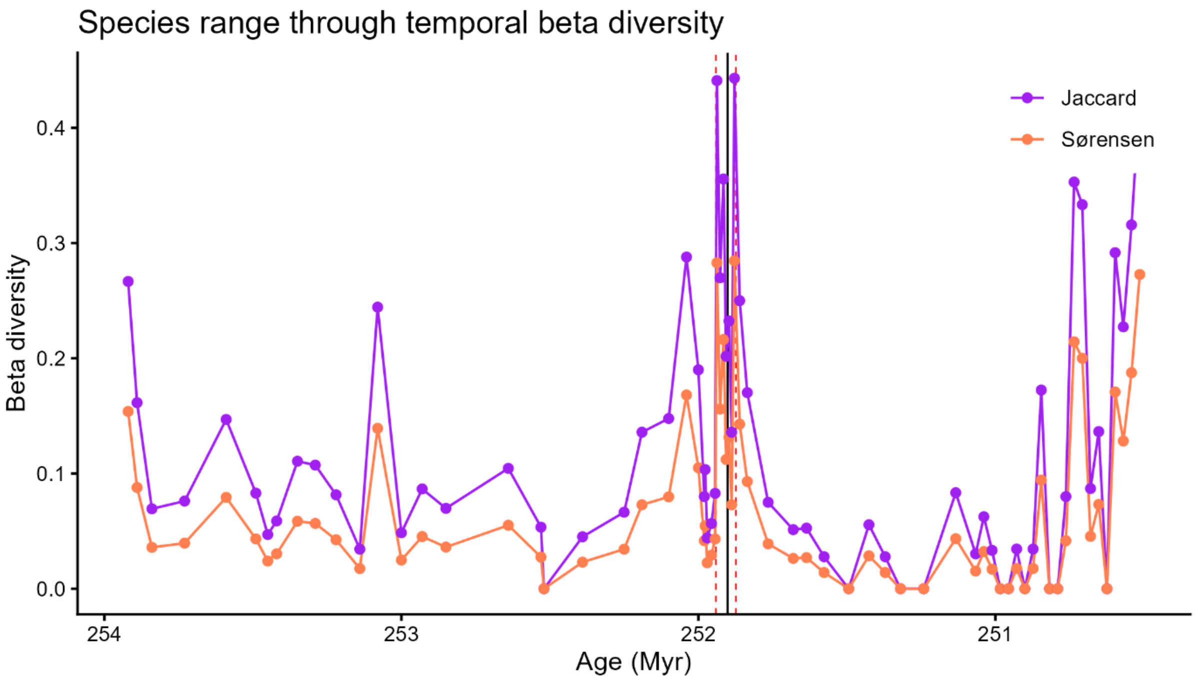


**Figure S5:** Jaccard and Sørensen pairwise dissimilarity indices between communities in two consecutive time bins. Red dashed lines indicate the onset and end of the extinction interval. Black vertical line indicates the Permian–Triassic boundary.
